# Rationalizing the Effects of RNA Modifications on Protein Interactions

**DOI:** 10.1101/2024.08.31.610603

**Authors:** Andrea Vandelli, Laura Broglia, Alexandros Armaos, Riccardo Delli Ponti, Gian Gaetano Tartaglia

## Abstract

RNA modifications play a crucial role in regulating gene expression by altering RNA structure and modulating interactions with RNA-binding proteins (RBPs). In this study, we explore the impact of specific RNA chemical modifications—N6-methyladenosine (m⁶A), A-to-I editing, and pseudouridine (Ψ)—on RNA secondary structure and protein-RNA interactions. Utilizing genome-wide data, including RNA secondary structure predictions and protein-RNA interaction datasets, we classify proteins into distinct categories based on their binding behaviors: modification-specific and structure-independent, or modification-promiscuous and structure-dependent. For instance, m⁶A readers like YTHDF2 exhibit modification-specific and structure-independent binding, consistently attaching to m⁶A regardless of structural changes. Conversely, proteins such as U2AF2 display modification-promiscuous and structure-dependent behavior, altering their binding preferences in response to structural changes induced by different modifications. A-to-I editing, which causes significant structural changes, typically reduces protein interactions, while Ψ enhances RNA structural stability, albeit with variable effects on protein binding. To better predict these interactions, we developed the catRAPID 2.0 RNA modifications algorithm, which forecasts the effects of RNA modifications on protein-RNA binding propensities. This algorithm serves as a valuable tool for researchers, enabling the prediction and analysis of RNA modifications’ impact on protein interactions, thus offering new insights into RNA biology and engineering. The catRAPID 2.0 RNA modifications tool is available at http://service.tartaglialab.com/new_submission/catrapid_omicsv2_rna_mod.

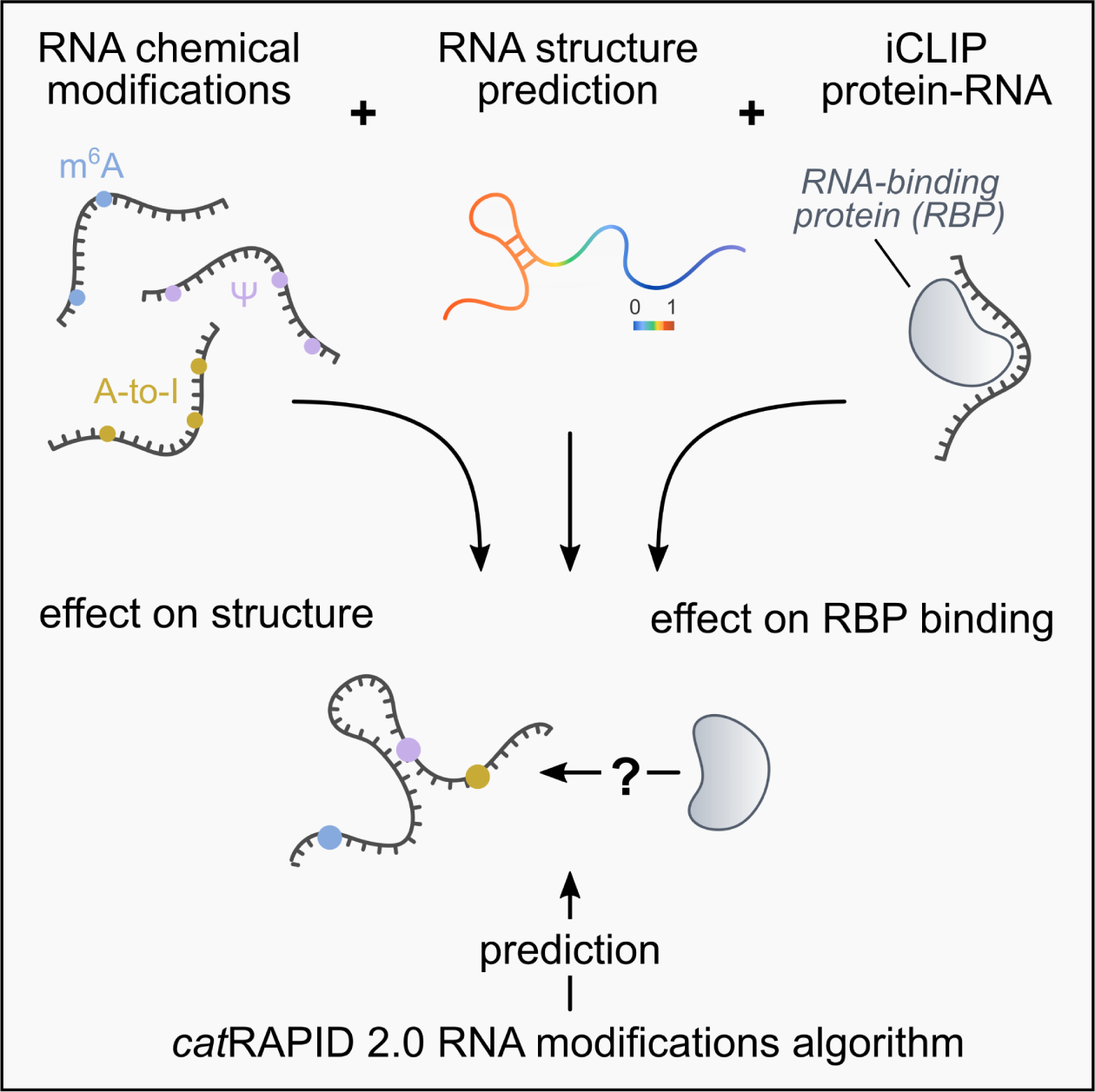

## Introduction

RNA molecules can undergo extensive chemical modifications, resulting in the formation of non-canonical nucleotides. These modifications are far from passive; they can significantly alter RNA reactivity by influencing base pairing, conformational dynamics and interactions with proteins, ultimately shaping the biological functions of the modified RNA. Modifications, such as N^6^-methyladenosine (m⁶A), N1-methyladenosine (m^1^A), 5-methylcytosine (m^5^C), and pseudouridine (Ψ), can change the chemical properties of RNA, affecting its structure, stability, and interactions with RNA-binding proteins (RBPs). For instance, m⁶A is the most prevalent modification in messenger RNAs (mRNAs) and non-coding RNAs, influencing RNA folding and the binding affinity of RBPs, thereby modulating gene expression ^1,2^. Similarly, m^1^A enhances RNA stability and translation efficiency ^3,4^ while Ψ promotes RNA base stacking, contributing to increased stability and improved translation efficiency ^5^.

By influencing RBP binding, RNA modifications regulate mRNA stability, splicing, and translation, thus controlling gene expression at multiple levels. These modifications also shape cellular responses to stress and other stimuli by modulating the assembly and function of ribonucleoprotein complexes, essential for processes such as splicing and translation ^5^. The study of RNA modifications, known as the epitranscriptome, is revealing new layers of gene regulation, emphasizing the importance of these chemical changes in maintaining cellular homeostasis and adapting to environmental challenges ^2,6^. Understanding these processes is vital for elucidating the complex regulatory networks that govern cellular function.

The m⁶A modification influences RNA structure and stability, primarily by reducing the double-stranded content of RNAs. Indeed, the methyl group in the adenosine renders the paring with U (m⁶A•U) less strong compared to the canonical A•U interaction^7^. Therefore, while m⁶A•U hampers the RNA capacity to adopt structured conformations, it favors the formation of unfolded and linear RNAs ^7^. The weakening of the Watson-Crick base pairing leads to changes in local RNA folding kinetics. These structural alterations significantly impact protein binding to RNAs, either facilitating or inhibiting specific interactions ^8,9^. For instance, heterogeneous nuclear ribonucleoprotein C (HNRNPC) preferably binds purine rich motifs that become linear and accessible upon m⁶A modification in the proximity regions ^9^.

In contrast, Ψ enhances RNA structural stability, increasing the double-stranded content within cells. Ψ promotes local RNA base stacking in both single- and double-stranded conformations, contributing to greater stability and rigidity of the RNA backbone. This increased stability is due to Ψ unique properties, which allow it to stack better than uridine (U) and to enhance neighboring nucleotide interactions ^10,11^. The major groove created by the RNA backbone upon Ψ modification becomes increasingly accessible for polar interactions with proteins ^11,12^. Yet, the Ψ of short structured RNAs decreases the flexibility of the RNA by increasing base stacking, consequently reducing the propensity of proteins such as MBNL1 to bind ^13^. In general, incorporating pseudouridine into mRNA improves translation efficiency by reducing the activation of RNA-dependent protein kinase (PKR), which otherwise inhibits translation. mRNAs containing pseudouridine instead of uridine exhibit reduced association with PKR, rendering the pseudouridine-containing mRNAs more efficiently translated ^14^. This suggests that pseudouridine not only stabilizes RNA structurally but also enhances its functional capabilities in cellular contexts, potentially increasing double-stranded RNA content ^14,15^.

A-to-I RNA editing exhibits a dual role in modulating double-stranded RNA (dsRNA) content, depending on the cellular context ^16–18^. Indeed, while inosine has a destabilizing effect on perfectly matched dsRNA duplexes, its presence serves to stabilize imperfect dsRNA regions ^16^. A-to-I editing can decrease dsRNA content by destabilizing existing dsRNA structures in specific contexts. Editing can alter the local RNA structure, affecting duplex stability and the accessibility of *trans*-acting factors ^19,20^. Even a single A-to-I change can impact RNA structure, as seen in editing-dependent structural changes in 3’ UTRs that reduce the accessibility of AGO2-miRNA complexes to target sites, thereby stabilizing the mRNA ^17^. Conversely, A-to-I editing can increase dsRNA content by creating new dsRNA structures from adjacent inverted repeats in the transcriptome. This is achieved through the editing of the pseudo-pairing A•C, generating an I:C pair that exhibits greater thermodynamic stability ^16^. A high number of A-to-I editing sites have been identified in non-coding regions, such as introns and 3′ UTRs, which harbor Alu retroelements ^21,22^. Perfect double-stranded regions formed by Alu repeats are subject to editing, resulting in a destabilizing effect on their structure ^16^. This structural rearrangement contributes to the MDA5 dsRNA sensing pathway, helping to prevent excessive activation of the immune response ^16,23^. In summary, A-to-I editing has a context-dependent effect on dsRNA content: it can generate new dsRNA structures from inverted repeats while also disrupting existing dsRNA by altering the local structure. The overall impact depends on the specific sequences and cellular conditions.

In this manuscript, we investigate the effects of RNA chemical modifications—specifically m⁶A, A-to-I editing, and Ψ —on RNA secondary structure and protein-RNA interactions. We identify the sites of these modifications and cross-reference them with iCLIP data to analyze how chemical modifications affect protein binding sites. Our findings reveal that m⁶A-modified RNAs attract more protein binders, A-to-I editing has the most pronounced structural impact, and pseudouridylation generally stabilizes RNA structures. By integrating changes in RNA secondary structure into the *cat*RAPID algorithm, we provide a foundation for understanding how chemical modifications influence RNA-protein interactions. To fully understand RNA modifications, it is essential to examine both the structural and chemical effects of these changes in combination with their impact on protein recruitment. This comprehensive understanding can inform therapeutic interventions not only in disorders involving alterations of the epitranscriptome but also in utilizing RNA modifications as tools, such as in RNA-based therapeutics. For further exploration, the catRAPID tool can be reached at http://service.tartaglialab.com/new_submission/catrapid_omicsv2_rna_mod.

## Results

To compare the effect of RNA modifications on the secondary structure, we first selected genome-wide experiments for three of the most studied RNA modifications: m⁶A, A-to-I, and Ψ ^24–26^. The datasets have different dimensions, with A-to-I including many more modified nucleotides (46286) compared to m⁶A (15188) and Ψ (1488). We used the *RNAfold* module from *Vienna* ^27^ to study the effect of the different modification on the RNA secondary structure (see **Materials and Methods**). For this analysis, we selected fragments of different sizes (50, 100, 200 nucleotides) centered on the modifications, and computed the RNA secondary structure for all the fragments with and without the corresponding modification in the central nucleotide (for *RNAfold* encoding: A=6 for m⁶A, A=I for A-to-I, U=P for Ψ). As expected, the effect of RNA modifications is not completely altering the RNA structure, with the majority of the fragments retaining the same structure (structural identity with/without the modification = 100%; **Figure 1A**). A-to-I is the modification with the strongest effect on the RNA structure, with 60% of the RNA structures altered by the modification (**Figure 1A**, **Supplementary Table 1**). By contrast, m⁶A and Ψ show a weaker effect on the RNA structure, with only few structures changing between with and without the modification. However, approximately 30% of the fragments in our dataset show significant alterations in secondary structure, with a structural identity of less than 75% (**Supplementary Table 1**). We note that 50 nucleotides fragments are the limit for the *cat*RAPID algorithm ^28,29^; however, this size may be too small to capture significant changes. Thus, 50-nucleotide fragments might not provide enough information on the RNA structure. In contrast, fragments of 200 nucleotides show marginal changes in their structure compared to fragments of 100 nucleotides (**Supplementary Table 1,2**). For this reason, as a good compromise between size and retained information, we used RNA fragments of 100 nucleotides for the following analysis.

**Figure 1:**
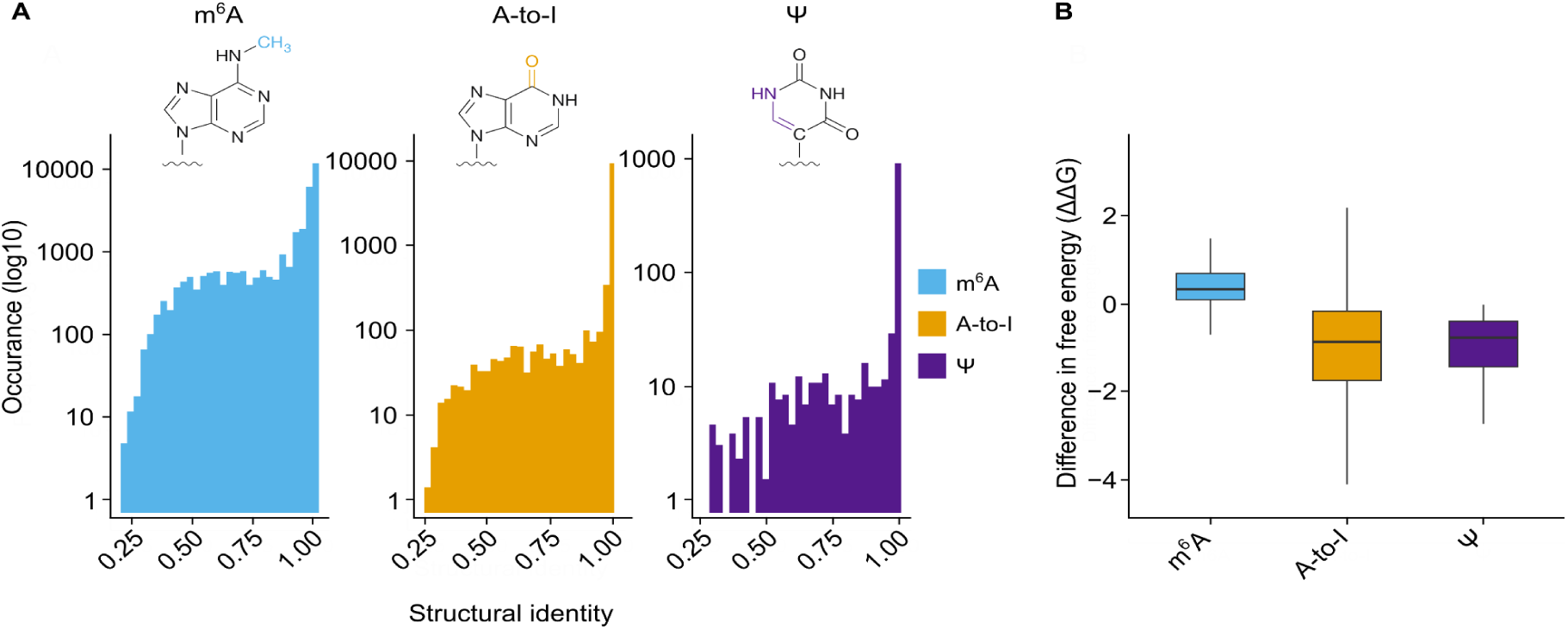
Impact of RNA modifications on secondary structure stability. The figure illustrates the distribution of RNA fragments based on changes in secondary structure upon modification with m⁶A, A-to-I editing, and pseudouridine (Ψ). **(A)** The percentage of RNA fragments that retain or alter their secondary structure after modification is depicted. A-to-I editing causes the most significant structural disruptions, while m⁶A and Ψ generally result in less pronounced changes. **(B)** The differences in RNA free energy (ΔΔG) between modified and unmodified fragments are shown, with positive ΔΔG values indicating decreased stability and negative values indicating increased stability. m⁶A leads to a loss of structural stability, aligning with its role in promoting single-stranded regions. Ψ, conversely, enhances RNA stability by decreasing ΔΔG, reinforcing its stabilizing effect on RNA structure.

To identify the RNAs that undergo the most significant structural changes due to a modification, we focused on fragments that exhibited the greatest change in structure following the modification (i.e., those with lower structural identity). We then compared these **structurally variable** fragments (i.e., structural identity < 100%) with the **structurally stable** fragments (i.e., structural identity = 100%). The most variable fragments are enriched in the regulation of CD8-positive, alpha-beta T cell differentiation, and lymphocyte homeostasis. These enrichments suggest that RNA modifications may play a critical role in immune responses by influencing RNA structure and its function in gene expression and cellular processes ^5^. Notably, many RNAs that become structurally destabilized after A-to-I editing are known to encode proteins involved in the innate immune response, particularly those linked to the type-I interferon pathway ^16^. This suggests that a delicate balance must be maintained between the levels of double-stranded RNAs within cells and the activation of the MDA5-dependent interferon response. Chemical modifications of RNA may be crucial in maintaining this balance, helping to regulate the interaction between RNA structures and the immune system’s signaling pathways.

To further understand the impact of RNA modifications on RNA structure, we analyzed the free energies of fragments whose structures were altered by the modification (i.e., structural identity < 100%). For each RNA fragment, we calculated the difference in free energy (ΔΔG) between the modified and unmodified structures using *RNAfold* ^27^. An increase in ΔΔG indicates that the structure has become less stable due to the modification, while a decrease in ΔΔG suggests that the structure has become more stable. Among the three modifications, m⁶A was the most disruptive to RNA secondary structure, with a ΔΔG > 0 in the majority of cases (**Figure 1B**).

This result is expected, as it is well known that m⁶A has a helicase-like effect on RNA secondary structure, promoting the formation of single-stranded nucleotides ^30^. In contrast, Ψ exhibits the opposite trend, decreasing the ΔΔG of RNA secondary structures. As observed in mRNA vaccine optimization, pseudouridylated RNAs are well-tolerated by cells, which improves the *in vivo* stability of modified RNA molecules ^31,32^. Additionally, the presence of Ψ stabilizes RNA duplexes, explaining why our results show that RNA structures are stabilized by this modification ^33^. A-to-I, on the other hand, shows a wider range of effects on RNA secondary structure, with case-specific instances where the modification can either stabilize or disrupt the RNA structure. This variability has been partially noted in previous literature, as there is still no general consensus on the effect of A-to-I on RNA structure. For example, some studies have found that A-to-I can both disrupt and promote double-stranded nucleotides ^34^.

Depending on the length of the RNA fragments, multiple modifications can also be present in the same region. While the central nucleotide of each fragment is always modified by design in our dataset, we do not have information about the epigenetic state of the neighboring nucleotides. For all the fragments in our dataset, we examined how many additional modifications were present. Interestingly, secondary structures that were not altered by the central modification (structural identity = 100%) tended to have fewer or no other modifications in nearby regions. In contrast, fragments that exhibited changes in their secondary structure (structural identity < 100%) generally had at least one additional adjacent modification on average (**Supplementary** Figure 2). On the one hand, these results might indicate that RNA modifications in close proximity within the same RNA molecule could have cumulative effects on RNA structure, leading to more significant structural rearrangements. It is known that RNA modifications *in cis* can exert a synergistic effect on functional outcomes ^35^. For instance, m^1^A and m^5^A act in concert to promote RNA degradation by favoring the interaction between the target RNA and the HSRP12-YTHDF2 complex ^36^. On the other hand, the structural change upon the RNA modification could enhance the exposure of regions previously inaccessible to other modifying enzymes. For example, the addition of m⁶A by METTL3/METTL14 in the 3’ UTR of p21 facilitates the nearby placement of m^5^C by NSUN2, and *vice versa*, synergistically promoting the p21 mRNA translation ^37^. The mechanism of this cross-talk is unknown, but the proximity of the methylated sites suggests that a structural switch could participate in the modifying enzyme recruitment.

### iCLIP-based analysis of protein interactions with modified RNAs

The next step in our work was to study the interplay between RNA modifications, secondary structure alterations, and interactions with proteins. In the previous paragraph, we showed how RNA modifications can have a different effect on the RNA secondary structure (**Figure 1**). To study how this affects the protein-RNA interactions, we collected iCLIP data from the POSTAR3 database (**Materials and Methods**) ^38^. By overlapping the coordinates of the peaks with those of the modified RNAs, we analyzed the average number of proteins binding to each modified RNA. Interestingly, A-to-I, which is the most prevalent modification in our dataset in terms of RNA fragments, is associated with the fewest protein binders per RNA (**Supplementary** Figure 2). In contrast, m⁶A-modified RNAs tend to be bound by a higher number of proteins.

In addition, we conducted a more protein-centric analysis, focusing on the number of interactions each iCLIP protein had with the modified RNAs. m⁶A stood out as the modification with the most protein interactions, showing approximately 13,600 interaction peaks (**Figure 2**).

**Figure 2:**
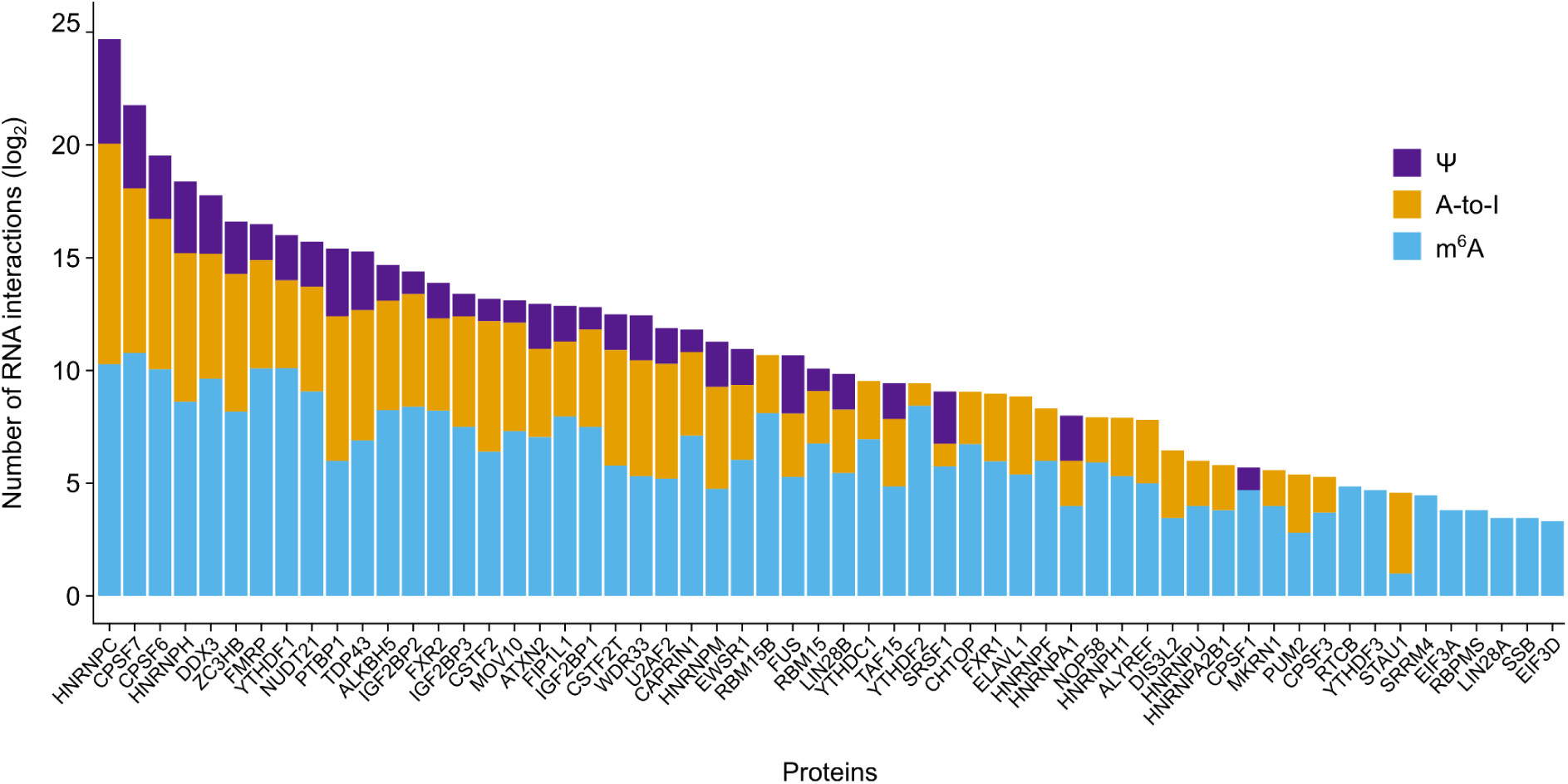
Protein interactions with modified RNAs as identified through iCLIP data. The total number of protein interaction peaks with RNA fragments containing specific chemical modifications—m⁶A, A-to-I editing, and pseudouridine (Ψ)—is presented. Bars represent the interaction frequency for each modification, emphasizing the differential binding preferences of proteins. m⁶A-modified RNAs display the highest number of protein interactions, indicating a strong affinity for m⁶A sites. In contrast, despite the higher occurrence of A-to-I modifications, these fragments exhibit fewer protein interactions, suggesting that the structural alterations caused by A-to-I editing may limit protein binding.

To further study protein preferences for RNA modifications, we calculated the total percentage of interactions each protein has with the three modifications in our dataset. Proteins that are highly-specific to a single modification exhibit 100% of their interactions with that modification, while proteins with broader specificity or modification-independent binding show varying percentages of interactions with the three different modifications. Moreover, proteins can be classified as either **structurally-dependent** or **-independent** based on whether their binding preference changes when RNA fragments undergo alterations in secondary structure (**Figure 3A** and **B**). The majority of the iCLIP proteins tends to specifically bind m⁶A, especially when the secondary structure is not altered by the modification (**Figure 4A**). However, structurally-variable RNA fragments exhibit a lower degree of binding for m⁶A, with an increment in binding fragments associated to A-to-I or Ψ. This result suggests how changes in RNA secondary structure affect the propensity of proteins binding a specific modification. In fact, when the RNA secondary structure is altered by the RNA modification, compared to the unmodified condition, proteins lose their preference for m⁶A and increase their bindings to other modifications.

**Figure 3:**
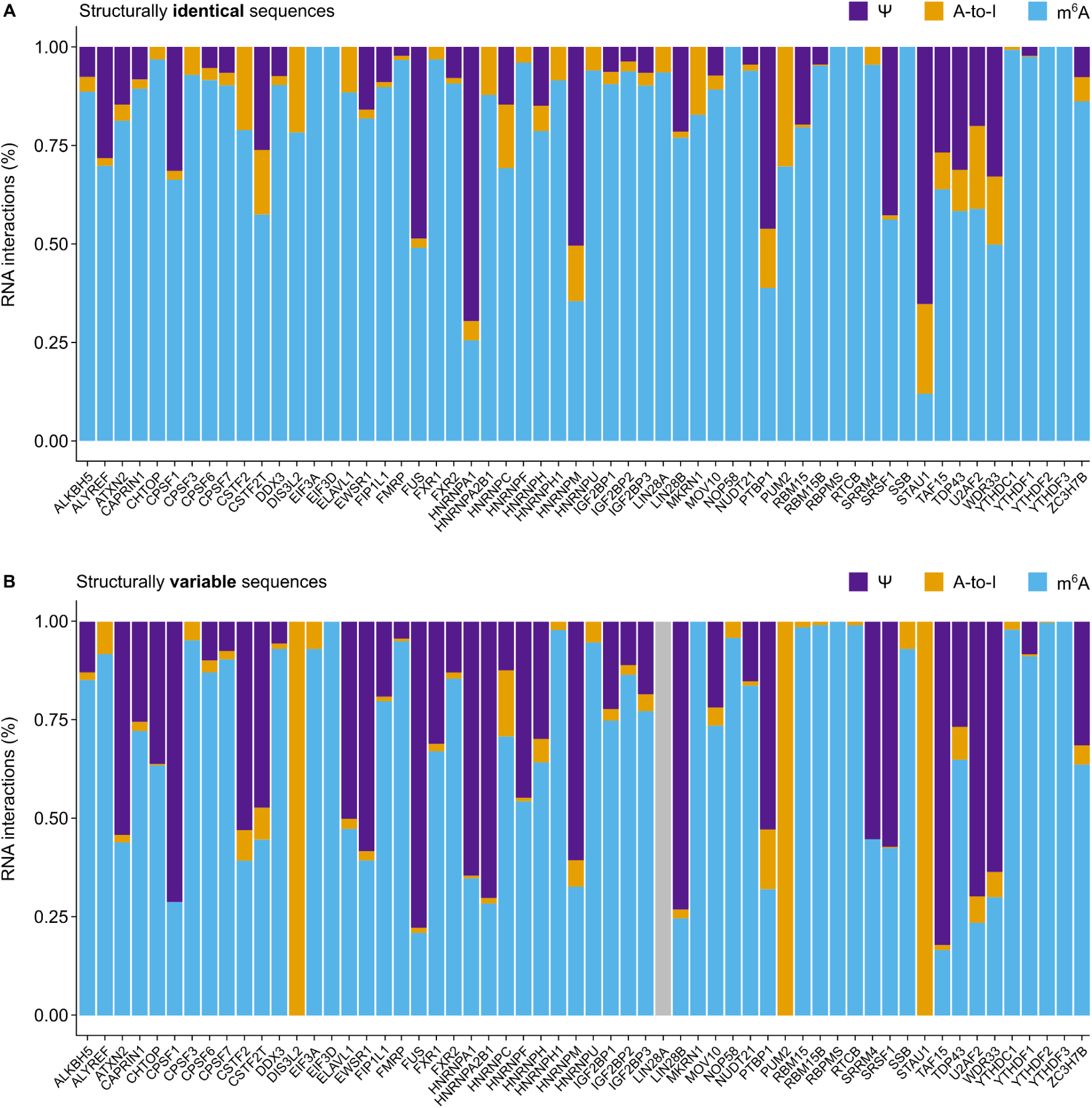
Protein binding specificity relative to RNA modifications and structural changes. Protein interactions with m⁶A, A-to-I, and Ψ modifications are displayed, focusing on both structurally stable and variable RNA fragments. **(A)** The bar plot illustrates the percentage of protein interactions specific to each modification, normalized by the number of structurally stable sequences, highlighting proteins that predominantly bind to a single modification, such as m⁶A. **(B)** The analysis of structurally variable sequences reveals how structural changes influence protein binding preferences. Proteins are shown to either maintain consistent binding to a specific modification regardless of structural alterations or shift their binding preferences in response to RNA structural changes.. Proteins without any RNA binder are highlighted in gray.

**Figure 4:**
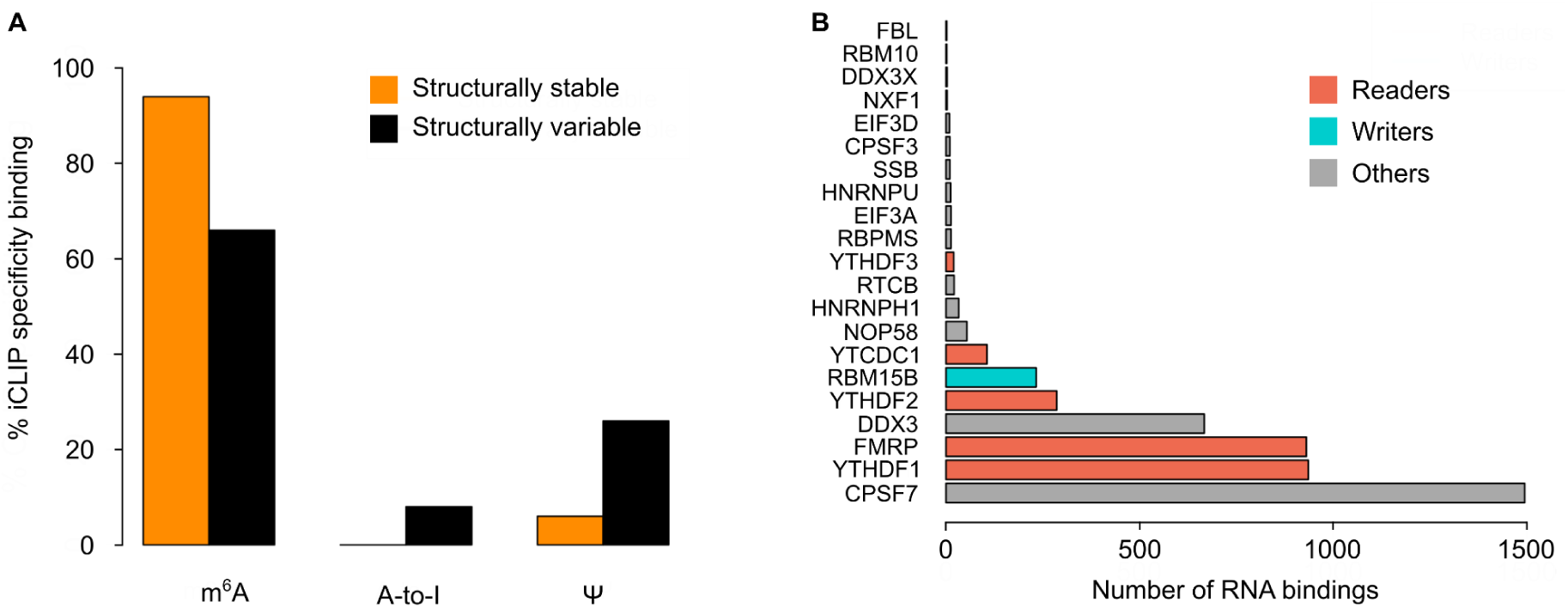
Protein binding specificity for each modification (iCLIP). (**A**) Barplot showing the modification preferentially bound (the maximum percentage of binding sites among the three modifications) by the iCLIP proteins. m⁶A is the modification preferentially bound by the majority of the proteins, but in structurally variable fragments this binding is significantly reduced. (**B**) Modification-specific and structure-independent proteins or proteins binding in >90% of the cases fragments with a specific modification without any preference for the structural stability of the sequence. Interestingly, three m⁶A readers and a writer appear among the list of 18 proteins. The absolute number of bindings with m⁶A structurally stable fragments for each protein is reported in the figure.

The binding percentage to each modification for each protein (or modification specificity) derived from the previous analysis can be used to further classify the proteins for their structure or modification specificity. Proteins changing their modification specificity for structurally stable or variable fragments are considered **modification-promiscuous** and **structure-dependent** (19 proteins; **Supplementary** Figure 3). Moreover, we can identify proteins that not only change their modification specificity between structurally stable and structurally variable fragments but also bind with higher specificity to the modification associated with the variable fragments (i.e., the maximum percentage of RNA bindings in stable fragments is smaller than the maximum percentage of RNA bindings in variable fragment; 9 proteins: PUM2, DIS3L2, TAF15, FUS, WDR33, U2AF2, SRSF1, STAU1, CPSF1). Although very little literature has reported on the effects of chemical modifications on protein binding, it has been shown that both m⁶A and Ψ modulate the binding of PUM2 ^39^ and U2AF2 ^40^.

Proteins binding in >90% of the cases a specific modification, both in structurally stable and structurally variable fragments, are defined as **modification-specific** and **structure-independent** (18 proteins; **Figure 4B**). If we select proteins always binding >90% of m⁶A fragments, in the proteins with the overall highest number of bindings with m⁶A structurally stable fragments, we identify 3 m⁶A readers (YTHDF1, YTHDF2, FMRP) and a writer (RBM15B; **Figure 4B**). Indeed, the m⁶A modification acts as a direct recognition element for m⁶A readers. YTH family m⁶A readers possess an aromatic pocket within their YTH domain that accommodates the modified nucleotide, irrespective of the surrounding sequence context. None of these YTH domains, except YTHDC1, exhibit sequence selectivity at the position preceding the m⁶A modification ^41,42^. FMRP has been demonstrated to preferentially bind m⁶A-modified mRNA, thereby negatively regulating their translation ^43^. The modification-specific and structure-independent protein category includes CPSF7, a factor involved in the regulation of pre-mRNA 3′ end processing by promoting proximal alternative polyadenylation and shortening of the mRNA 3′ UTRs ^44^. A recent study has demonstrated that CPSF7 binds to 2′-O-methylated transcripts, thereby promoting alternative polyadenylation ^45^. Given that m⁶A modifications are enriched near stop codons and 3′ UTRs ^25,46^, CPSF7 may also be recruited by m⁶A modifications to regulate 3′ processing, as it is enriched in m⁶A-modified regions ^47,48^. Among the proteins that bind to m⁶A-modified RNAs, we identified DDX3, a DEAD-box RNA helicase known to interact with the m⁶A RNA demethylase ALKBH5, highlighting a potential role for DDX3 in m⁶A regulation ^49^. Helicase activity appears to play a role in m⁶A function, as evidenced by the presence of a helicase domain belonging to the DEAD Box family in the m⁶A reader YTHDC2 ^50^. Additionally, DDX5, another DEAD-box helicase, acts as a cofactor in the m⁶A regulatory network by interacting with YTHDC1 and controlling circular RNA biogenesis ^51^.

After identifying m⁶A readers as the most modification-specific and structure-independent proteins for m⁶A, we decided to further investigate how these readers bind and their relationship with changes in RNA structure. In this context, m⁶A serves as an ideal case study, since m⁶A readers, writers, and erasers have been extensively studied ^52^. By analyzing the iCLIP data, m⁶A readers appear to be the proteins with highest preference to bind m⁶A modified fragments, compared to their total number of iCLIP peaks (**Supplementary** Figure 4).

However, different readers exhibit varying preferences for binding to m⁶A-modified fragments. This result suggests the presence of a wide set of readers with different specificities. In general, m⁶A readers tend to bind more to structurally stable fragments (2746 stable bound RNAs vs 470 variable bound RNAs), even though this could be partially due to the proportion of structurally stable and variable m⁶A sequences (13015 stable fragments vs 2173 variable fragments). Interestingly, HNRNPC is the reader with the least specificity for m⁶A fragments (**Supplementary** Figure 4).This could be attributed to HNRNPC unique binding mechanism, which targets structural changes near the methylated nucleotides rather than recognizing the methylated residues themselves ^9^. This mechanism likely broadens its target specificity to a wider range of RNA fragments.

Given the varying specificities of m⁶A readers (**Supplementary** Figure 4), we hypothesized that structural changes in RNA might influence their binding preferences. To explore this, we calculated the difference in the number of single-stranded nucleotides for each m⁶A fragment bound by these readers, comparing the modified and unmodified conditions (**Figure 5**).The analysis revealed that some readers preferentially bind to RNAs that become more linear upon m⁶A modification, while others are more attracted to sequences that become more structured. This modification leads to a significant shift in the binding affinity of various RNA-binding proteins towards single-stranded RNA. Specifically, ELAVL1, FMRP, and HNRNPC exhibit a higher affinity for single-stranded RNA regions compared to YTHDC2 and HNRNPA2B1.

**Figure 5:**
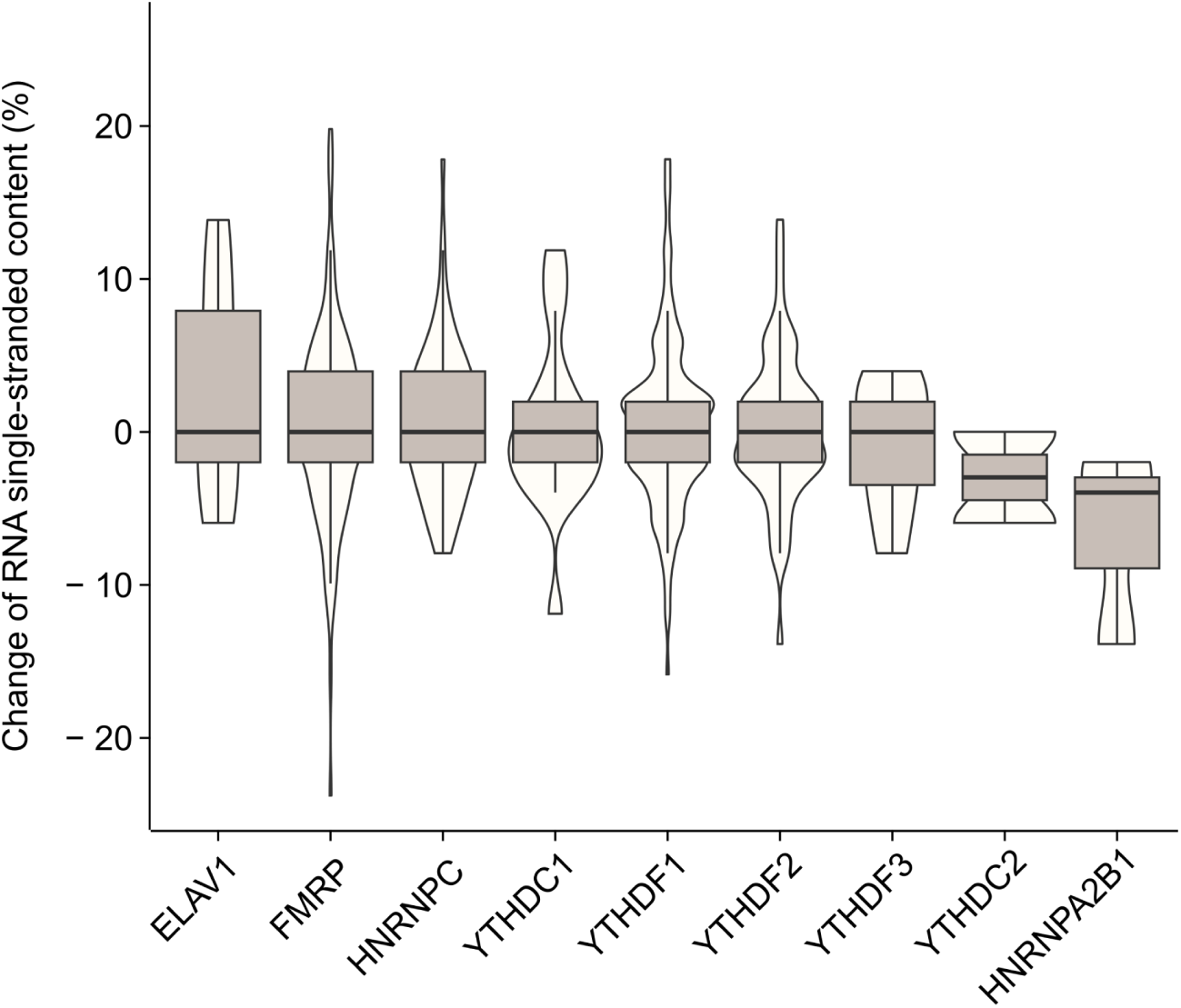
Binding preferences of m⁶A readers in response to RNA structural changes. Changes in single-stranded content of RNA fragments are shown, comparing the RNA structure before and after m⁶A modification. The y-axis quantifies the percentage change in single-stranded regions, with positive values indicating increased linearization and negative values indicating a shift toward more structured conformations. Certain m⁶A readers, such as ELAVL1 and HNRNPC, display a preference for RNA regions that become more linear following m⁶A modification. In contrast, other readers, like YTHDC2, are drawn to regions that acquire additional structure.

HNRNPC, for instance, shows a preference for purine-rich motifs that become linear and more accessible after m⁶A modification, which enhances its binding to these regions ^9,48^ . Similarly, the presence of m⁶A may alter RNA structure in a way that increases FMRP’s ^53^ and ELAVL1’s accessibility to RNA, especially near m⁶A-binding motifs ^54,55^ .

In contrast, HNRNPA2B1, despite its ability to bind m⁶A at both linear and structured RNA sites, does not show the same preference for single-stranded RNA as HNRNPC and ELAVL1. This is likely due to its capacity to bind m⁶A in both unstructured/linear regions and more complex structured regions through its multiple RNA recognition motifs ^56^. YTHDC2, on the other hand, relies more on non-m⁶A-dependent interactions with U-rich motifs via its various RNA-binding domains (R3H, helicase domain, and OB fold) and is less influenced by m⁶A modifications, leading to reduced binding to single-stranded RNA regions modified by m⁶A ^57–59^.

### Rationalizing protein-RNA binding effects of RNA modifications using catRAPID

iCLIP experiments provide reliable information on protein binding preferences. However, only 79 proteins have been analyzed using iCLIP, and the varying experimental conditions make it difficult to directly compare the results across studies. Predictive algorithms such as *catRAPID* (**Materials and Methods**) were developed to provide new insights on a vast number of protein-RNA interactions, especially in cases where experimental data are lacking ^28,29^. In several analyses we used this algorithm to successfully predict interactions to study RNA viruses ^60–63^, long non-coding RNAs ^64,65^ and phase-separating condensates ^66,67^.

While RNA modification data is becoming increasingly available, there is still limited knowledge about how these modifications influence protein-RNA interactions. In order to fill this gap, we developed *catRAPID 2.0 RNA modifications*, an algorithm to predict how the protein-RNA binding is affected by the RNA modifications. The algorithm is based on *RNAfold* to predict the secondary structure upon modification ^27^, and the original *cat*RAPID 2.0 *omics* algorithm ^29^ to predict RNA-protein interactions. The input RNA can have one or multiple modifications chosen among pseudouridine, m⁶A and inosine, that have to be coded in the sequence as P, 6, I, respectively. The interaction propensity score between the molecules is then calculated both with and without RNA modification, and the difference between the two interaction propensities (ΔI) is the result for each protein-RNA pair. A very high or very low ΔI indicates a strong difference in binding propensity upon modification. The algorithm also reports the difference in RNA free energy (or ΔΔG) to highlight the change in RNA secondary structure upon modification.

To validate the *cat*RAPID 2.0 *RNA modifications* algorithm, we predicted the interactions between the human RBPome and the RNA fragments (**Materials and Methods**), generating a total of over 109 predicted interactions. We then focused on the 79 proteins for which we also had experimental data, allowing us to compare the algorithm’s predictions with real-world observations and assess its accuracy.

For each RNA-binding protein (RBP), we gathered its iCLIP interactions with RNA fragments that include the specific chemical modification, labeling these as “positive” interactions. These were then compared to a control group, consisting of an equal number of “negative” sequences—1,000 randomly selected RNA fragments that also contain the chemical modification but do not show any interaction with iCLIP peaks. This comparison allowed us to assess the specificity and strength of the protein-RNA binding influenced by the modifications. We predicted both positive and negative interactions using *cat*RAPID *2.0 RNA modifications*, calculating their binding propensity scores with and without the modification. For each protein, we compute the ‘Target recognition ability,’ which is a Z-score calculated to determine how often the positive iCLIP interactions has higher scores compared to the randomly extracted negatives (**Materials and Methods**). Our analysis reveals that the change in interaction propensity upon modification is significantly higher for the positive interactions than for the negatives (**Figure 6**). This analysis uses the maximal interaction within the iCLIP data as a representative value for each RBP, although it yields similar results when using the average value.

**Figure 6:**
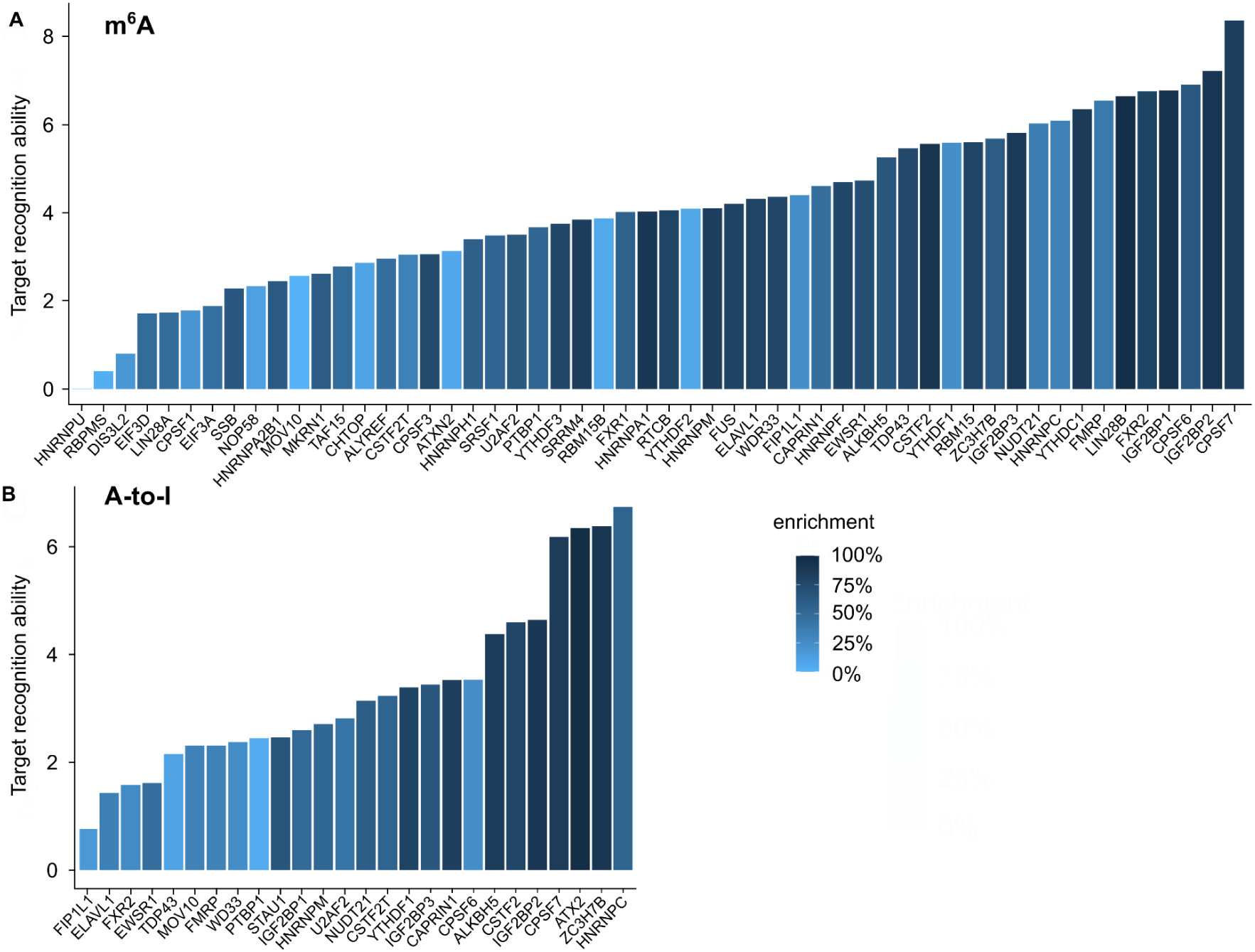
Influence of RNA modifications on protein-RNA interaction predictions. The impact of m⁶A and A-to-I modifications on the binding propensity of proteins to RNA is depicted through Z-scores (called ‘Target recognition ability’), calculated using the catRAPID 2.0 RNA modifications algorithm. **(A)** Z-scores represent the effect of m⁶A on interaction propensity, with higher scores indicating a stronger modification-induced change in binding. Bars are color-coded to reflect enrichment levels, which indicate how frequently positive iCLIP interactions score higher than randomly selected negative sequences. **(B)** A similar analysis is shown for A-to-I modifications, demonstrating a variable impact on protein-RNA binding. The results reveal that specific proteins exhibit significant changes in binding affinity upon modification, highlighting the predictive power of the algorithm and providing insights into how RNA modifications can alter protein-RNA interactions.

To ensure meaningful statistics, only proteins interacting with at least 10 modified RNA fragments were considered for each modification. The presence of the m⁶A modification led to a 75% enrichment in the Z-score compared to control interactions, observed in over 30 proteins of interest (**Figure 6A**). Similarly, for A-to-I modifications, 10 proteins exhibited an enrichment of over 75% (**Figure 6B**). There is a significant correlation between the Z-score and enrichment for both m⁶A (r = 0.56, P-value < 1.18e-05) and A-to-I (r = 0.73, P-value < 2.608e-05). However, for Ψ, meaningful results could not be extrapolated due to the presence of only two proteins with more than 10 interactions.

## Conclusions

In this study we aimed to rationalize the effects of RNA chemical modifications on protein interactions. We build on an approach previously introduced to study mutations affecting protein aggregation ^68,69^. We utilized genome-wide data on RNA modifications, secondary structure predictions, and protein-RNA interaction datasets to elucidate the roles of these chemical changes.

At the structural level, we found that while most RNA fragments with m⁶A show minor alterations, a significant subset (∼30%) display considerable changes associated with reduced stability, consistent with m⁶A’s role in promoting single-stranded regions. ^7,70^. A-to-I has the most pronounced impact, with 60% of modified fragments undergoing substantial structural changes, reflecting its complex role in RNA editing and regulation ^3,16^. Ψ generally stabilizes RNA structures, supporting its role in enhancing RNA stability and function *in vivo*^33^.

To understand the role of chemical modifications in protein binding, we combined genome-wide data on RNA modifications ^24–26^ with iCLIP data from different labs, all consistently using the HEK293 cell line. While maintaining the same cell line across studies provides some level of consistency, it is important to acknowledge that differences in experimental setups and conditions could influence the results. Factors such as variations in experimental protocols, reagent quality, and environmental conditions under which the cells were maintained can all contribute to discrepancies and variations in data outcomes. Despite these potential limitations, we adopted rigorous methods to ensure the reliability and relevance of our findings. By carefully selecting high-quality datasets and standardizing the analysis procedures, we aimed to minimize the impact of these variables. For this purpose, we collected protein-RNA interactions from the POSTAR3 database ^38^.

We found that m⁶A-modified RNAs attract more protein binders compared to other modifications. This modification showed for instance strong interactions with known m⁶A writers such as YTHDC1. Indeed, the binding affinity of several proteins to m⁶A-modified RNAs is influenced by the RNA’s structural context, with notable shifts in binding partners when the RNA structure is altered by the modification ^2^. A-to-I-modified RNAs, on the other hand, showed fewer protein interactions, likely due to the modification’s disruptive effects on RNA structure ^71^. Ψ generally has a stabilizing influence on RNA structure, but its effects on protein-RNA interactions can vary depending on the context ^72^. Indeed, although Ψ is often associated with enhanced RNA stability, this does not always translate to increased interactions with RBPs. For instance, the addition of Ψ within the binding motif UGUAR for PUM2 reduces binding affinity ^39^ . Similarly, when uridines are substituted with Ψ within CUG repeats, the RNA becomes less flexible, which reduces the binding of MBNL1 ^13^.

Based on our analysis, RBPs have been classified based on their binding specificity and dependence on RNA structure. Modification-specific and structure-independent proteins predominantly bind to m⁶A-modified RNAs, whereas modification-promiscuous proteins exhibit variable preferences depending on the RNA structural changes.

To study how RNA modifications affect protein binding in general, we modified *cat*RAPID, a computational algorithm designed to predict the binding propensity between RNA and proteins ^28^, to create our newly developed *cat*RAPID 2.0 RNA modifications algorithm. This enhanced tool provides predictive insights into how RNA modifications influence protein-RNA binding propensities, expanding the scope of studying RNA-protein dynamics by modeling interactions involving multiple modifications. The *cat*RAPID algorithm combines information about RNA secondary structure, hydrogen bonding, and van der Waals interactions to estimate how likely a specific RNA-protein pair is to interact ^64^. The major hypothesis behind *cat*RAPID 2.0 RNA modifications is that the main impact of chemical modifications on RNA is structural. This structural alteration hypothesis is based on our analysis showing significant changes in RNA stability and conformation upon modification. However, it is important to acknowledge that other factors beyond structural changes are also relevant for the interaction between modified RNA and proteins. These factors include the intrinsic physico-chemical properties of the modifications themselves. For instance, modifications can alter the hydrophobicity, charge, and steric properties of nucleotides, which can directly affect how RNA interacts with various proteins ^73^. The binding affinity and specificity of RBPs may be influenced by these physico-chemical changes, independent of structural alterations. Examples of such physico-chemical influences include electrostatic interactions, where modifications can change the charge distribution on the RNA, influencing how it interacts with positively or negatively charged domains of RBPs. For instance, in N1-methyladenosine (m^1^A), the methylation occurs in the Watson-Crick interface, giving rise to the formation of a positively charged base that gives the possibility for strong electrostatic interactions with proteins ^3,12^. In other cases, the addition of a methyl group at the carbon-5 position of cytosine, forming 5-methylcytosine (m^5^C), can also modify RNA hydrophobicity. This methylation enhances base stacking between adjacent nucleotides and increases the hydrophobicity of the RNA major groove, thereby facilitating hydrophobic interactions. ^12,74^. Thus, changing the RNA chemical properties such as the introduction of hydrophobic groups, can affect the solubility and aggregation properties of RNA, altering its interactions with hydrophobic or hydrophilic protein surfaces ^73^. Moreover, modifications can introduce bulky groups that may either facilitate or obstruct protein binding sites on the RNA, depending on the spatial configuration ^75^. These factors are not explicitly accounted for in our hypothesis. Nevertheless, they play a crucial role in determining the full spectrum of RNA-protein interactions. Future research should aim to integrate these physico-chemical aspects with structural data to provide a more comprehensive understanding of how RNA modifications influence molecular interactions. Such a holistic approach will enable us to better elucidate the mechanisms by which RNA modifications regulate gene expression and other cellular processes.

There are several potential avenues for future research that could further illuminate the roles of RNA modifications and their broader implications.

Firstly, expanding the dataset to include more RNA modifications beyond m⁶A, A-to-I, and Ψ would provide a more comprehensive understanding of how different chemical changes affect RNA structure and function. Modifications such as m^1^A, m^5^C, and others could be investigated to determine their unique structural impacts and their influence on protein binding ^76^. Additionally, leveraging high-throughput sequencing technologies and novel experimental techniques could identify more modification sites and provide higher resolution data on RNA-protein interactions. Advancements in high-throughput technologies will open the possibility to not focus only on one modification at the time, but to look at the interplay of different RNA chemical modifications and how their combined effect alters RNA-protein interactions.

Secondly, the dynamic nature of RNA modifications in response to cellular signals and environmental changes warrants further exploration. Investigating how these modifications are regulated in different cellular contexts, such as during stress responses, development, or disease states, could reveal critical insights into their functional roles ^77^. Single-cell RNA sequencing combined with modification-specific detection methods could elucidate how modifications contribute to cellular heterogeneity and plasticity ^78^.

Thirdly, translating these findings into therapeutic applications offers a promising future direction. RNA modifications have significant potential in therapeutic development, particularly in the design of more stable and efficient mRNA vaccines and therapeutics. Understanding the mechanisms by which modifications affect RNA stability and protein interactions could lead to the development of novel RNA-based treatments for various diseases, including cancer, viral infections, and genetic disorders. Additionally, targeting the enzymes responsible for adding or removing RNA modifications presents a potential strategy for modulating gene expression and treating diseases associated with dysregulated RNA modification patterns.

## Materials and methods

### RNA datasets

We selected the genomics coordinates of the three RNA modifications from genome-wide experiments on human cells ^24–26^. We selected these experiments to have the same cell lines (HEK293/HEK293T) and genome version (hg19) for the three modifications. For m⁶A, the data provided a window of different sizes, so we selected the central coordinate as representative for m⁶A modifications. For A-to-I, the data provided results for three replicates, so we selected coordinates in common between the replicates as representative for A-to-I modifications. For Ψ we didn’t have genomic coordinates but the position on the reference transcript. For this case, we used all the exon coordinates for each gene and mapped the reference position on the transcripts to obtain the corresponding genomic coordinates.

For each genomic coordinate, we extracted the corresponding fragment by using *BEDTools getfasta* ^79^. After that, we selected the closest corresponding nucleotide for that specific modification (A for m⁶A and A-to-I; T/U for Ψ) and opened a window around it based on desired fragment length (± 25 nt for fragments of length 50, for example). Fragments including undefined nucleotides (N) were removed from the dataset.

### Secondary structure predictions

We used the command line version of *RNAfold* to compute the RNA secondary structure of all the fragments in our dataset, with standard settings ^27^. We computed the RNA secondary structure both for the WT fragments and for the modified ones. To add the modification, we used the reported alphabet for *RNAfold* to accordingly modify the central nucleotide in the fragments: A=6 for m⁶A, A=I for A-to-I, T/U=P for Ψ. After obtaining both the WT and modified RNA secondary structure, we compared the dots-and-brackets profiles to obtain the structural identity.

### Protein-RNA interactions

We collected protein-RNA interactions (peaks from the data) obtained through iCLIP technique from POSTAR3 database, which stores protein-RNA binding sites found with different computational tools and techniques across various organisms ^38^. We selected only peaks coming from HEK293/ HEK293T cell lines and we mapped them to the human genome (h19 version) using the tool CrossMap ^80^ in order to be consistent with the RNA dataset. The peaks coming from different replicates and with different tools were merged using the BEDTools ‘merge’ utility. Then, only merged peaks overlapping across all computational tools for binding site identification were collected and among those we filtered out those not supported by a minimum number of replicates, following a workflow adopted in a previous publication ^78,81^. After the filtering step we ended up with a total of 79 proteins and ∼500.000 peaks.

### *cat*RAPID RNA modifications validation

*cat*RAPID is an algorithm designed to estimate the binding propensity of protein-RNA pairs by integrating factors such as RNA secondary structure, hydrogen bonding, and van der Waals interactions. It effectively distinguishes between interacting and non-interacting pairs, achieving an area under the ROC curve of 0.78 based on training with approximately half a million experimentally validated interactions ^29,82^.

To enhance the accuracy of RNA secondary structure predictions, we updated catRAPID ^28^ by incorporating the latest version of *RNAfold*, which predicts the effects of chemical modifications ^27^, while maintaining the original algorithm’s structure without reparametrization. For its validation, we compared iCLIP interactions (positives) with randomly extracted sequences (negatives) using *cat*RAPID 2.0 RNA modifications to predict binding propensity scores with and without modifications. To quantify the impact of RNA modifications on protein-RNA interactions, we calculated a Z-score for each RBP, using the formula (|ΔI| - mean) / sd, where |ΔI| is the absolute difference between modified and wild-type interaction propensity scores, and the mean and sd are calculated from the positive and negative sets. Our analysis showed that positive interactions exhibited a significantly greater change in interaction propensity |ΔI| upon modification compared to negatives, validating the accuracy of our predictions. These calculations used the maximal interaction within the iCLIP data for each RBP, though similar results were observed when using the average value.

### *cat*RAPID RNA modifications webserver

The *cat*RAPID 2.0 RNA modifications module can predict protein-RNA interactions of RNA sequences containing modified residues selected from inosine (I), pseudouridine (P), dihydrouridine (D), m⁶A (6), 7DA (7), and nebularine (9). *cat*RAPID 2.0 RNA modifications can be used to compute the interactions between the human RBPome (2064 proteins) proteome and the modified RNA fragments (62962 sequences). This version of the algorithm is available as a webserver at (http://service.tartaglialab.com/new_submission/catrapid_omicsv2_rna_mod).

*cat*RAPID 2.0 RNA modifications takes as input a list of modified RNAs of interest and one of the 8 available proteomes and computes the differences in interactions propensities with both the wild-type and modified sequences, providing:

● ΔΔG: The difference in RNA energy due to the modification.
● ΔI: The difference in interaction propensity between the modified and unmodified states (ΔI = Ix - Iy).

## Data Availability Statement

The datasets generated and analyzed during the current study are available from the corresponding author on reasonable request. The catRAPID 2.0 RNA modifications algorithm, used in this study, is publicly accessible at http://service.tartaglialab.com/new_submission/catrapid_omicsv2_rna_mod. Data supporting the findings of this study, including a table with secondary structure predictions of all the fragments, are provided within the supplementary materials.

## Acknowledgements

The authors would like to thank the other members of Tartaglia’s group, especially Dr. Alessio Colantoni and Gabriele Proietti.

## Contributions

RDP and GGT conceived and designed the study. RDP, LB, AA and AV performed the analysis and assembled the figures. RDP, AV, LB, AA and GGT wrote the manuscript. All the authors read and agreed with the content of the manuscript. AA designed the webserver.

