## Supplementary Materials for "Rationalizing the Effects of RNA Modifications on Protein Interactions"

| | | Fragment lenghts | | | | | |
|---|---|---|---|---|---|---|---|
| | | 50 nt | | 100 nt | | 200 nt | |
| | | Structural identity <100% | Structural identity <75% | Structural identity <100% | Structural identity <75% | Structural identity <100% | Structural identity <75% |
| modifications | $m^6A$ | 1820 (12%) | 987 (6%) | 2173 (14%) | 959 (6%) | 2356 (15%) | 732 (5%) |
| | A-to-I | 28045 (60%) | 10614 (23%) | 20190 (60%) | 9670 (21%) | 9670 (64%) | 7480 (16%) |
| | Ψ | 262 (17%) | 156 (10%) | 294 (20%) | 154 (10%) | 312 (21%) | 131 (9%) |

**Supplementary Table 1**: Dataset composition of structurally variable fragments: the table shows the raw numbers and percentages of sequences with structural identity < 100% and < 75% for different lengths (50, 100, 200 nucleotides) and for the three target modifications ($m^6A$, A-to-I, Ψ).

**Supplementary Table 2:** We report the genomic location, sequence, and secondary structures of both unmodified and modified RNA sequences, along with the measured changes resulting from the three modifications—$m^6A$, A-to-I, and Ψ—considered in this study.

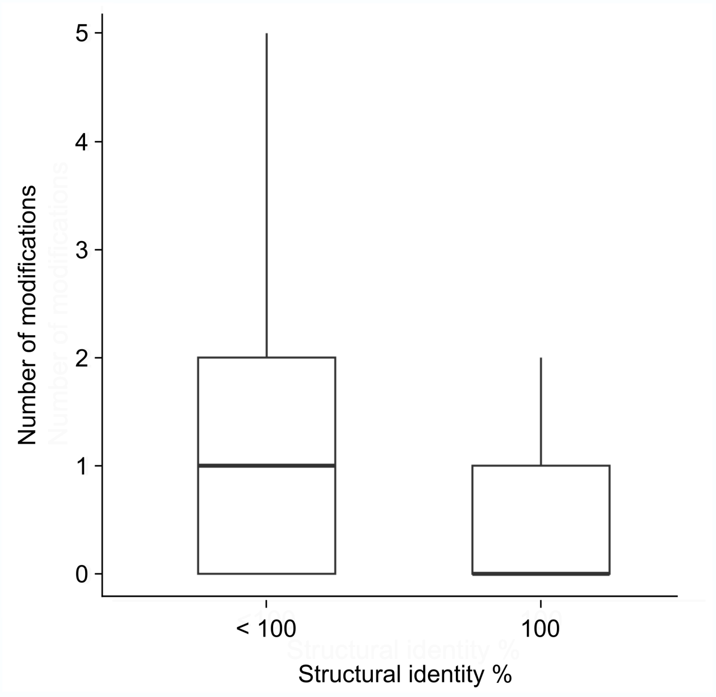


**Supplementary Figure 1** Number of adjacent modifications found in RNA fragments with stable and variable secondary structure. Variable fragments seem to be more prone to accept other modifications in the vicinity. The number of modifications doesn't include the central nucleotide, which is considered 0.

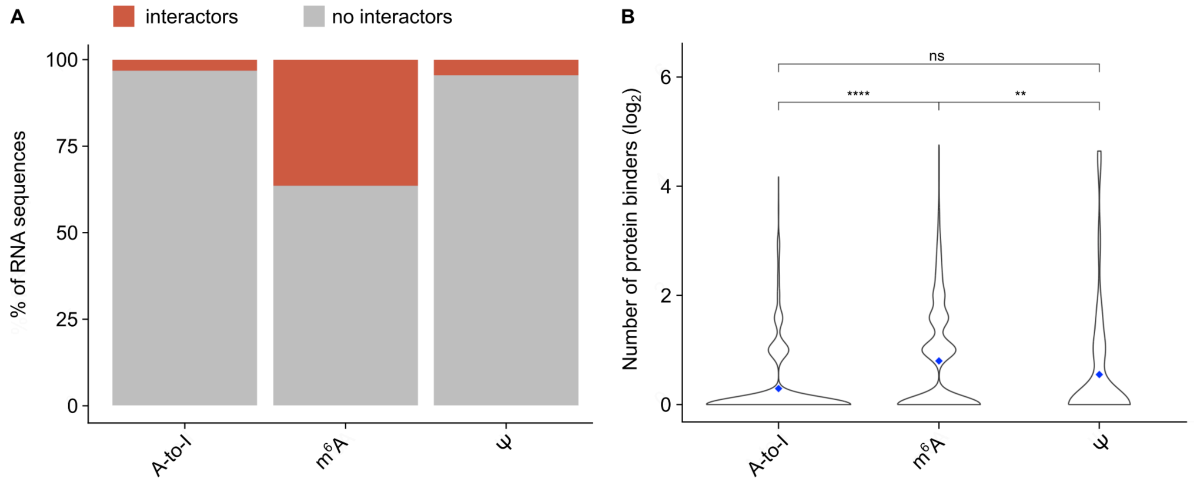


**Supplementary Fig. 2**: Number of different protein binders (iCLIP) for each modified RNA fragment. (**A**) Percentages of modified fragments with (orange) and without (gray) protein interactions, respectively. (**B**) Distribution of the number of protein binders for each modification ($\log_2$). The average number of interactors for each modification is shown as a blue square. In general, $m^6A$ shows the highest number of interactions per sequence, even though the A-to-I modification dataset contains a much higher number of fragments.

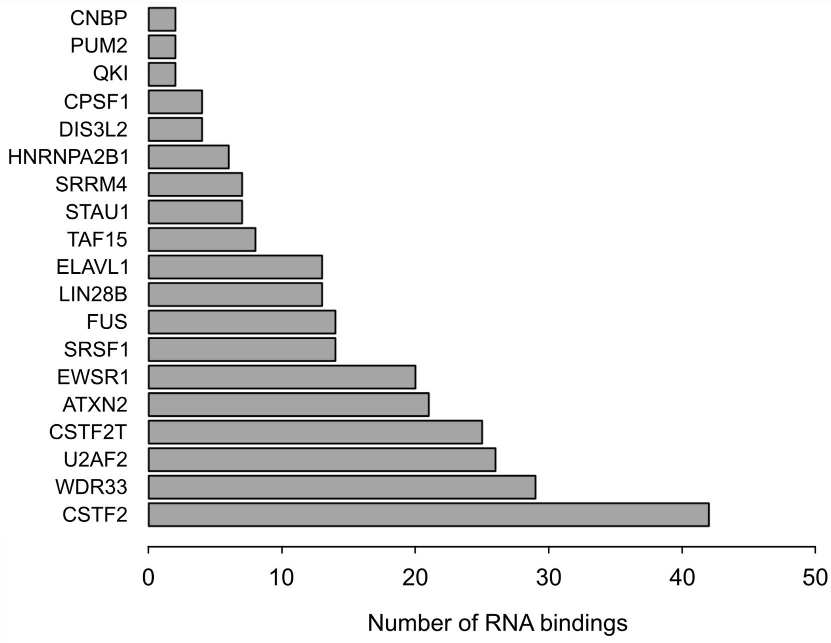


**Supplementary Figure 3:** The analysis identifies proteins that change their binding preferences between structurally stable and variable RNA fragments, particularly those showing a higher binding specificity to modifications in structurally variable contexts. The bar plot highlights proteins that demonstrate increased affinity for RNA fragments that undergo significant structural changes due to modifications. These proteins, including TAF15, U2AF2, and PTBP1, exhibit a shift in binding from $m^6A$-modified to Ψ-modified fragments, reflecting their dependency on RNA structural alterations. The absolute number of RNA bindings for each protein is reported, providing a quantitative view of how structural variability influences protein interaction with modified RNA.

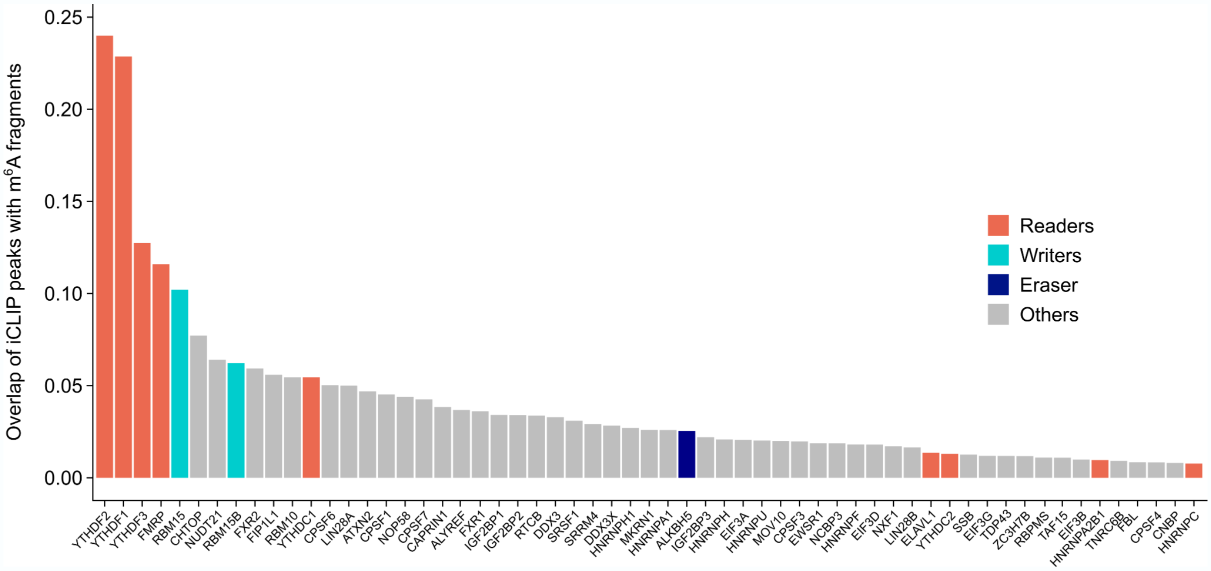


**Supplementary Figure 4. Differential binding preferences of m$^6$A readers, writers, and erasers.** We report the proportion of m$^6$A-modified RNA fragments bound by various proteins, categorized as readers, writers, and erasers of the m$^6$A modification. Each bar represents the fraction of total iCLIP peaks that correspond to m$^6$A-modified sites for individual proteins. The data reveal significant variation in binding preferences among m$^6$A readers, with proteins such as YTHDF1 and YTHDF2 showing a strong affinity for m$^6$A-modified RNA, while others, like HNRNPC, exhibit less specificity. This variation suggests that different m$^6$A readers may have distinct roles in recognizing and interacting with m$^6$A-modified RNA, depending on the structural context and the presence of additional RNA features.
